# Don’t Dismiss Logistic Regression: The Case for Sensible Extraction of Interactions in the Era of Machine Learning

**DOI:** 10.1101/2019.12.15.877134

**Authors:** Joshua J. Levy, A. James O’Malley

## Abstract

**Background:** Machine learning approaches have become increasingly popular modeling techniques, relying on data-driven heuristics to arrive at its solutions. Recent comparisons between these algorithms and traditional statistical modeling techniques have largely ignored the superiority gained by the former approaches due to involvement of model-building search algorithms. This has led to alignment of statistical and machine learning approaches with different types of problems and the under-development of procedures that combine their attributes. In this context, we hoped to understand the domains of applicability for each approach and to identify areas where a marriage between the two approaches is warranted. We then sought to develop a hybrid statistical-machine learning procedure with the best attributes of each.

**Methods:** We present three simple examples to illustrate when to use each modeling approach and posit a general framework for combining them into an enhanced logistic regression model building procedure that aids interpretation. We study 556 benchmark machine learning datasets to uncover when machine learning techniques outperformed rudimentary logistic regression models and so are potentially well-equipped to enhance them. We illustrate a software package, *InteractionTransformer*, which embeds logistic regression with advanced model building capacity by using machine learning algorithms to extract candidate interaction features from a random forest model for inclusion in the model. Finally, we apply our enhanced logistic regression analysis to two real-word biomedical examples, one where predictors vary linearly with the outcome and another with extensive second-order interactions.

**Results:** Preliminary statistical analysis demonstrated that across 556 benchmark datasets, the random forest approach significantly outperformed the logistic regression approach. We found a statistically significant increase in predictive performance when using hybrid procedures and greater clarity in the association with the outcome of terms acquired compared to directly interpreting the random forest output.

**Conclusions:** When a random forest model is closer to the true model, hybrid statistical-machine learning procedures can substantially enhance the performance of statistical procedures in an automated manner while preserving easy interpretation of the results. Such hybrid methods may help facilitate widespread adoption of machine learning techniques in the biomedical setting.

## Background

In the era of Big Data, models with highly complex specifications are employed to study nontrivial biomedical phenomena, including genetic or epigenetic interactions, high-resolution modalities such as Computed Tomography (CT) scans and histopathology slide images [1–6]. Many statistical approaches to modeling these complex data rely on expert consultation and annotation to help determine which variables and targets to study. Oddly, despite the advances in computing power, these procedures have not been augmented with sophisticated model-building algorithms. At best commercial software still only supports simple and restrictive search strategies such as forward, backward and stepwise selection. In contrast, machine learning techniques employ a data-driven set of heuristics to simultaneously build and estimate models that may include an extensive array of nonlinear interactions in the data. While machine learning methods are particularly helpful in exploratory or predictive settings, not much may be revealed about how the set of covariates vary with each other and the dependent variable. Unlike statistical models, which are able to make predictions when the data has a low number of features as compared to the number of training samples, machine learning algorithms are much more suitable for the high dimensional domain, especially if the data is subject to multicollinearity, and to the low dimensional domain when it is not clear how the covariates vary with the outcome. For instance, machine learning technologies have demonstrated the ability to make impressive predictions on medical images, genomic data and Electronic Health Record (EHR) modalities in the presence of many training instances [7–9]. In recent years, these approaches have gained much traction in the biomedical space and will continue to do so in the years to come.

However, with researchers and practitioners flocking to adopt these new technologies, there are concerns that traditional statistical methods are being passed over too quickly. Many published papers that show that machine learning techniques outperform traditional statistical models. Yet, many of these papers mislead the reader by presenting unfair comparisons between the methodologies, for instance selecting datasets that are difficult to learn given insufficient model-building strategies or by comparing a statistical procedure with no embedded learning component to machine learning procedures [10]. Meanwhile, these featured datasets run counter to more clinically focused sets on which one prior study has demonstrated no performance improvement for machine learning techniques [11]. Given this ambiguity and the clear need for models that are inherently interpretable for widespread adoption in the biomedical space, we believe there is much to be gained by entwining these methodologies rather than thinking of them as competitors.

We address a gap among traditional statistical procedures by proposing that sophisticated search algorithms be paired with them to suggest terms that can be added to the base specification to improve model fit and the predictive performance of traditional statistical models. These search algorithms would consider sets of candidate predictors from sets as expansive as those available to machine learning algorithms. Whereas the implementations of traditional statistical procedures such as linear and logistic regression in commercial packages often include basic search algorithms (e.g., forward, backward and stepwise regression), these search algorithms would span all forms of higher-order interactions and nonlinear transformations. Crucially, they would not be limited to only the models that can be formed by the predictors in the model specification but rather would include a myriad of additional predictors formed from functions of one or more of them and nonlinear transformations of their sum. Therefore, they would substantially enhance model-building capacity, potentially bringing the performance of traditional statistical methods on par with machine learning methods while retaining the easy and direct interpretability of the results of statistical procedures. In addition, such hybrid procedures would enjoy the automatic implementation and scalability enjoyed by the machine learning methods.

In this paper, we aim to illustrate the domains from which machine learning models can be sensibly applied to assist traditional statistical models from which to generate models that can be widely adopted in clinical applications.

First, we describe the machine learning and logistic regression models and their differences, including scenarios under which each outperforms the other. Then, we demonstrate that extracting interactions via the machine learning can enhance logistic regression (hybrid approach) as well as the ability of logistic regression to “protect the null hypothesis” by inhibiting the additional of unwarranted interaction terms to the model. Finally, we show consistencies between the logistic regression model and the machine learning model when the logistic regression model is closer to the true model. This study shows that machine learning can be deployed to assist with the development of statistical modeling approaches to obtain machine-learning level performance with statistical model interpretability.

### Review of Modeling Approaches

We now introduce hallmark models from machine learning and traditional statistics to better highlight domains from which either technique may excel.

Statistical Analysis System (SAS) is perhaps the most widely used commercial statistical software package. Its regression procedures are well established [12]. Some procedures, such as Proc reg and Proc logistic, include simple search algorithms that allow progress to be made towards a best fitting model. For example, forwards, backwards and stepwise regression can be implemented with the mere inclusion of an optional term in the specification of the procedure. Although these procedures have high utility and a long history (stepwise regression first made its debut in 1960), their model building search capacity has to our knowledge not been significantly enhanced since 1992 [13–15]. During this time, computational power has exponentiated. Remarkably, more expansive and otherwise advanced search algorithms have not been incorporated. Why not? Perhaps a lack of focus on, or accommodation of, predictive problems involving large data sets with many predictors. Whatever the reason, the computational power has for some time been available to enable these procedures to be enhanced. Therefore, we view this omission or gap as an historical anomaly that we seek to correct herein.

In the literature, comparisons are made between statistical models devoid of model-building and machine learning approaches enhanced by very sophisticated and powerful algorithms [10, 11]. This is an unfair comparison that risks making traditional statistical methods be dismissed too rapidly from consideration when choosing an analytic approach to a problem.

### Logistic Regression

Logistic regression is the most widely used modeling approach for binary outcomes in epidemiology and medicine [16]. The model is a part of the family of generalized linear models that explicitly models the relationship between the explanatory variable X and response variable Y. Conditional on the predictors, a binary outcome Y is assumed to follow a binomial distribution for which p=P(Y = 1 | X) represents the probability of the binary response given the predictors:

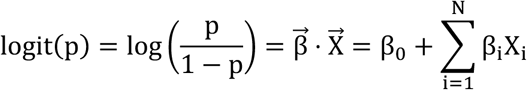

The approach above assumes a linear relationship between logarithm of the odds of the outcome and the predictors as equivalently depicted below:

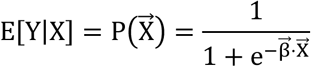

In applications involving a large number of predictors, LASSO or Ridge regression techniques serve to penalize the model’s complexity to make it more generalizable to unseen data by reducing the magnitude of the model coefficients such that high magnitude features become less tailored to the data used to tune the model parameters. Penalization also aids model estimation by repelling parameters from boundary values in much the same way that prior distributions with continuous support nudge the posterior distribution away from the extremities of the values a model parameter can attain.

### Classification and Regression Trees

Classification and Regression Trees (CART), otherwise known as Decision Trees [17], are a series of computational heuristics that arrive at a solution for a problem by forming binary splitting rules on the features of the data based on the criterion of maximizing the information gained about the outcome from making the split. This can be formulated mathematically as:

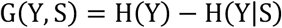

where S is an indicator variable indicating which side of the split contains a given observation. The above equation states that the information acquired about a set of labels given the split is the entropy of the original set of labels minus the entropy of the labels given the split, where entropy is defined as:

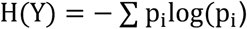

The entropy conditioned on the split is the weighted average of the entropy of the labels partitioned to each child, weighted by the number of samples sent to each child:

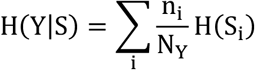

An alternative criterion that can be used in this framework, and the one used in this work, is Gini Impurity, which measures the probability of incorrect identification of a randomly chosen element and is given by:

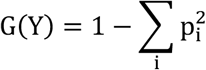

Decreases in Gini Impurity are expressed similarly to information gain, and both represent key criterion to decide which features and values should be used to split a branch. The values that the branches split on can be viewed as non-linear transforms of the data, while splits on multiple variables introduce interactions between those variables. A collection of hyperparameters effectively prune and limit the depth of each tree to generalize the model to unseen data.

### Random Forest

Random forest is an extension of the base CART algorithm by considering the construction of multiple CART models to arrive at a prediction [18, 19]. While decision trees are formed by utilizing all of the features, each CART model considers the best splitting feature out of n < N randomly selected features at a time, and fits subsequent branches by selecting out of another random set of n features, all while bootstrapping the training samples. A collection of these estimators is formed, and results are aggregated to form predictions. This is done because each of the CART models may provide high variance estimates by over-fitting their training observations. When aggregating results the variance is typically reduced by de-correlating the individual CART models by bootstrapping both the observations and the predictors. The number of trees and various aspects of each estimator’s construction can be limited in order to reduce overfitting the training data.

## Use Cases for Each Modeling Technique

Here, through a series of simple examples, we illustrate when either the traditional statistics or machine learning modeling approaches more closely resembles the true model and should be employed.

In figure 1a, we have a use case where we have a single linear continuous predictor. The distribution of the response is generated conditionally on this input by the true model:

**Figure 1:**
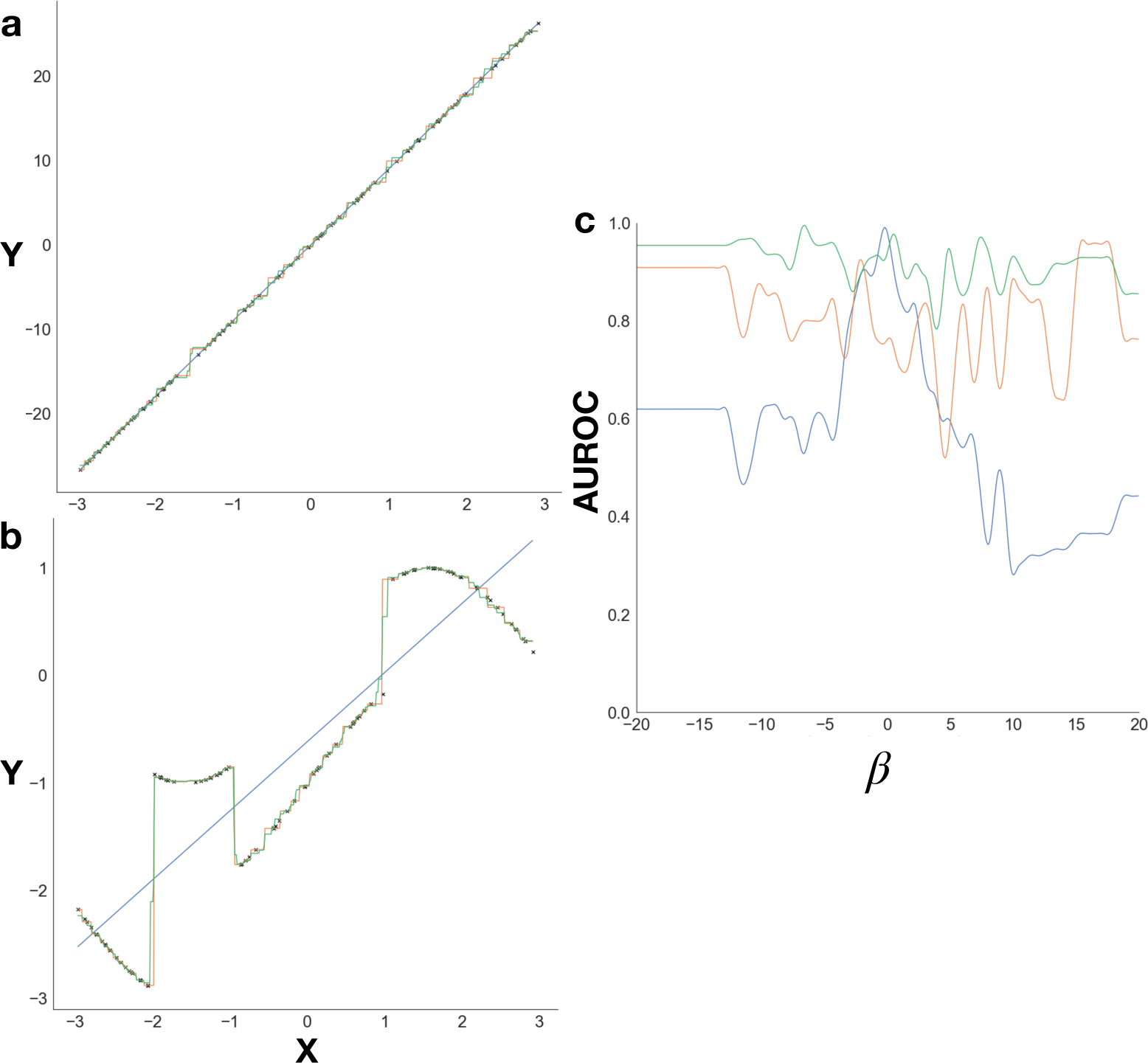
Characteristics of linear modeling approach (blue), decision trees (orange), random forest (green), original generated points (black); (a-b) Predictor X versus response Y for the a) linear continuous predictor, b) non-linear transformed predictor; c) binary prediction performance as a function of interaction strength, smoothed using Savitzky-Golay filter to best illustrate trends

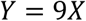

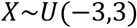

It seems upon inspection that a line passing through the datapoints could best describe this relationship, so first we fit a linear regression model to the data. The derived slope was found to be exactly 9 and the model had a mean absolute residual (MAR) of 0 on a held-out validation set. This example depicts the case when the true model is a linear regression model. Now, we fit two CART-models, a decision tree and the random forest model, and we see that the true linear relationship was approximated by these machine learning approaches using a staircase type fit. Under this model, the machine learning approach is guaranteed to have a higher residual error, and upon inspection of held out test data, this is the case (MAR of decision tree and random forest of 0.38 and 0.29 respectively). This illustration demonstrates that linear modeling approaches (any model of the generalized linear model family) are, in cases when a predictor varies linearly with the outcome, not made inadmissible by the random forest and other ML approaches. This means that we should be cognizant of cases when the true model space is linear because simpler models produced via this assumption may lay closer to the true model than a heuristic that only forms a coarse approximation.

Conversely, we designed two simple examples to illustrate when the machine learning approach is closer to the true model. In one example, figure 1b, we transform the continuous predictor X into the dependent outcome variable Y via the model:

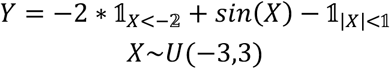

These transformations are difficult to be approximated by models that suppose the predictors are linearly associated with the outcome. We see that the linear fit cuts through the middle of the entire dataset and as such has a worse goodness of fit (mean absolute residual of 0.46) than the machine learning approaches (0.07 and 0.05 for decision tree and random forest respectively). This is because CART techniques do quite well in capturing discontinuities in the data and are able to come up with sharp and variable decision curves. In this case, the machine learning model better captures transformations to the data than the linear modeling approach.

As the last use case, figure 1c, we now consider a binary response model with four continuous predictors. The predictors vary linearly with the outcome; however, in the true model, the final two predictors interact:

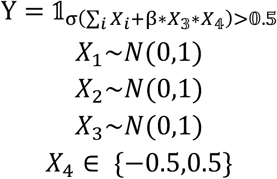

where:

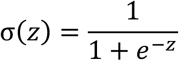

For this experiment, we vary the strength of the interaction β, and test the logistic regression, decision tree and random forest models, hoping to find that the logistic regression model performs worse as the magnitude of the interaction increases. This would mean that without adding interaction terms to the logistic model, the approach would be unable to detect the presence of two interacting predictors. In figure 1c, we see that the accuracy of the linear model diminishes in the domain of high interaction strength, while outperforming the machine learning approaches when the true model discards interactions and the relationship between predictors and response are linear. It can be noted, that the CART models’ accuracy does not significantly differ across varying interaction strengths.

In some cases, we can make reasonable guesses at which interactions would be the most appropriate to include such that the logistic model can better approximate the interaction relationship. While we had demonstrated the superior performance of statistical models when a linear solution space exists, when this solution space becomes more complex, the machine learning based approaches are better able to efficiently search over numerous combinations and specifications of the predictors separately and in combination. Traditional statistical models are not embedded with these machine learning model building features. These two points lead naturally to consideration of procedures that use machine learning methods to enhance traditional models by bolstering their model building capacity.

### Proposed New Method

Our proposed method utilizes machine learning to suggest terms to add to traditional statistical models; we refer to this as our hybrid statistical inference approach. We consider this approach a candidate to move on from the stepwise regression approaches currently enabled in SAS Proc Reg and SAS Proc logistic. We propose to augment the current SAS procedures with the output from a sophisticated and comprehensive search algorithm in the form of an ordered list of candidate second-order interactions. Linear and logistic regression procedures can then subsequently be estimated with the most promising of these terms included as predictors. While we illustrate our idea using this particular case, we also advocate for the development of far more expansive algorithms and thus types of inputs to further enhance traditional statistical methods. Such procedures may be characterized by a need to iterate between traditional statistical methods and machine learning methods such that each informs the improvement made to the other in the next iteration.

The hybrid approach stems from principles of explainable machine learning. Notably, many machine learning approaches struggle to gain traction and widespread adoption because they are unable to explain why they made their prediction. Existing techniques to explain the output from random forest algorithms attempt to quantify the marginal association of a variable’s capacity to reduce the amount of mislabeling in its decision splits. In contrast, the coefficients of the logistic regression model can be exponentiated to form an odds-ratio. However, these methods find global importance for the importance of each predictor and not the importance of the predictor for each test instance. The ideal interpretation technique for a machine learning model should project the random forest fit onto a linear subspace to determine the overall extent to which a specific factor contributes to the fit. Our hybrid approach utilizes such a projection to efficiently derive important model additions.

## Methods

Here, we discuss sensible approaches to enhance statistical modeling. Fundamentally, the essence of our approach is to build an intuitive understanding of when the “statistical” model (e.g. linear regression, logistic regression) is more representative of the true model that describes the data compared to the machine learning approach and what to do in cases where there is extensive incongruence. When the machine learning model is closer to the true approach, we posit that its output contains keys in the form of interactions between predictors and nonlinear transformations of predictors that can be exploited to substantially improve the fit of the corresponding statistical model. That is, machine learning models can be used as an efficient way of traversing the space of higher-order and nonlinear predictors to identify the best candidates to improve the fit of statistical models beyond that possible with main variable predictors alone. We conjecture that the addition of a relatively small number of these more complex forms of predictors will almost always lead to an improvement in fit of the statistical model to such an extent that its performance is similar to that of the best machine learning models. The greater ease of interpretability of statistical models then makes them highly competitive alternatives to machine learning alone. Inspired by a work that used these concepts to derive meaningful variable transformations [20], and as a special case of our proposed new methodology, we extend these ideas to automate the extraction of meaningful interactions via machine learning that are then added to the statistical modeling technique to enhance its performance. We also provide user-friendly software to allow users to apply this methodology to their own studies.

### SHAP Interaction Terms Extend Linear Model Building Capabilities

A nice feature of ML is that a number of algorithms have been developed that are amenable to hybridization with traditional statistical procedures or that provide the ingredients for making such a hybrid procedure. For example, SHAP (SHapley Additive ExPlanations) [21] is an algorithm designed to help interpret why any model has made its prediction for individual testing instance. These approaches explain “black-box” models by fitting linear additive surrogate models to each testing sample. If machine learning models can be characterized by complex curves over multiple variables, these models can be thought of as local tangent lines near each combination of variables present in the dataset (locally accurate). The coefficients of these models, dubbed shapley values, represent the importance of each predictor to the outcome prediction; these coefficients directly sum to the model prediction (additive), and features that are important to the prediction over a number of samples are consistent. SHAP sums the contributions of each feature to the model’s prediction over all possible permutations of other features.

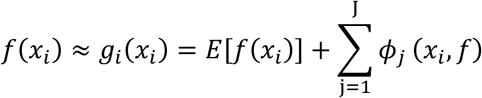

### Interaction Extraction and Autonomous Application

If the logistic regression model is correct, whether or not it contains interactions, then it will outperform any other specification. If it is not the correct model, then an approach with powerful search capability over the predictor space is likely to find ways to outperform logistic regression. If quantified in terms of functions of predictors and fed into logistic regression, these identified ways of improving model fit have the potential to enhance the fit of the logistic regression model such that its performance is more in-line with that of a search algorithm. For example, a subsequent logistic regression model augmented with complex predictors recommended by the output of a machine learning algorithm might then be sufficiently close to the truth that it outperforms all competitors. If not, one can continue iterating between the procedures until superior performance (and convergence) is obtained.

While the Surrogate Assisted Feature Extraction (SAFE) Transformer [20] has demonstrated the ability to accurately characterize variable transformations by estimating how a nonlinear predictor varies with the outcome, recent advances in SHAP methodology have allowed for the accurate detection of salient interactions of tree-based models [22, 23]. Simply put, these methodologies expand the aforementioned SHAP method to include interaction terms for individual samples. A measure of the global importance of these interactions can be characterized by summing the interaction effects over all of the samples to find the salient interactions for the model’s prediction. These important interactions can be extracted and added as predictors to the logistic regression model. A related operation is the use of machine learning algorithms to identify nonlinear transformations of variables that can be used to form other additional predictors. This is outside the scope of the current study, which is solely focused on the identification the candidate interaction effects and to provide advice of when they should be included.

Although interaction extraction has been studied previously, very few studies have attempted to characterize the importance of extracting these interactions over many datasets, and the ones that have focused on selecting a small number of predictors from which to test for interactions [4, 24]. Our novel contribution is the development of an automated means from which to extract these interactions and use them to enhance statistical model-based analyses of datasets of interest.

### Software Implementation and Availability

We have bundled and wrapped our multi-component methodology into a publicly available open source software package, InteractionTransformer, (GitHub: https://github.com/jlevy44/InteractionTransformer) that can be run using both R (GitHub via r-devtools: jlevy44/*interactiontransformer*) and Python (PyPI: *interactiontransformer*) to make these methods easily deployable for the biomedical researcher. InteractionTransformer takes as input the design matrices and outcome variable and fits a specified machine learning model to the data. Then, it extracts the most important SHAP-derived interaction terms from the machine learning model and augments the design matrices of the training and test data to include these interaction terms. It also provides additional capabilities to plot the salient SHAP features of any model and extract the variable importance assigned to all of the interaction terms for additional inspection. A wiki page detailing installation and usage can be found here: https://github.com/jlevy44/InteractionTransformer/wiki.

### EVALUATION OF HYBRID APPROACH: Dataset Acquisition and Experimental Description

We seek to estimate the general utility of extracting these salient interactions and when they should be sensibly employed by characterizing their activity over a large dataset.

We first acquired a dataset that would allow us to establish the necessary domains for the application of these modeling techniques. Inspired by a meta-analysis of random forest versus logistic regression, we utilized data from the OpenML database [25] to extract 556 datasets purposed for binary classification problems. Each dataset was cleaned and preprocessed using a data pipeline implemented in python.

We ran five-fold cross validation of naive implementations of L2-regularized logistic regression and random forest models using the scikit-learn and imbalanced-learn packages [26, 27] to compare the overall accuracy of the random forest models versus logistic regression models. Then, we characterize the extent to which extracting these interactions improve the fit of the logistic regression model. Since the primary domain of research pertinent to this study involves fully identified models, we excluded high dimensional genomics datasets and retained 277 datasets in the p<n domain; i.e., the number of observations always outnumbers the number of predictors. Then, we extracted the pertinent interactions from the training data of each of these datasets using *InteractionTransformer*. After applying these interaction terms to the design matrices of the training and test sets, we scored a naive logistic regression model using five-fold cross-validation to compare with the original two models. This will characterize the overall extent to which these interactions enhance the logistic regression model.

## Results

### Random Forest Outperforms Logistic Regression

A total of 556 OpenML datasets were fitted and tested on both the naive logistic regression and random forest models. Five-fold cross validation scores of the C-statistics (Area Under the Receiver Operating Curve, AUROC) of the models demonstrated that random forest models clearly outperform the logistic regression models by 0.061 AUROC on average (Figure 2a) (t=13.6, p=2.3e-36), where the random forest models scored 0.87 and logistic regression models scored an average cross-validated C-statistic of 0.81.

**Figure 2:**
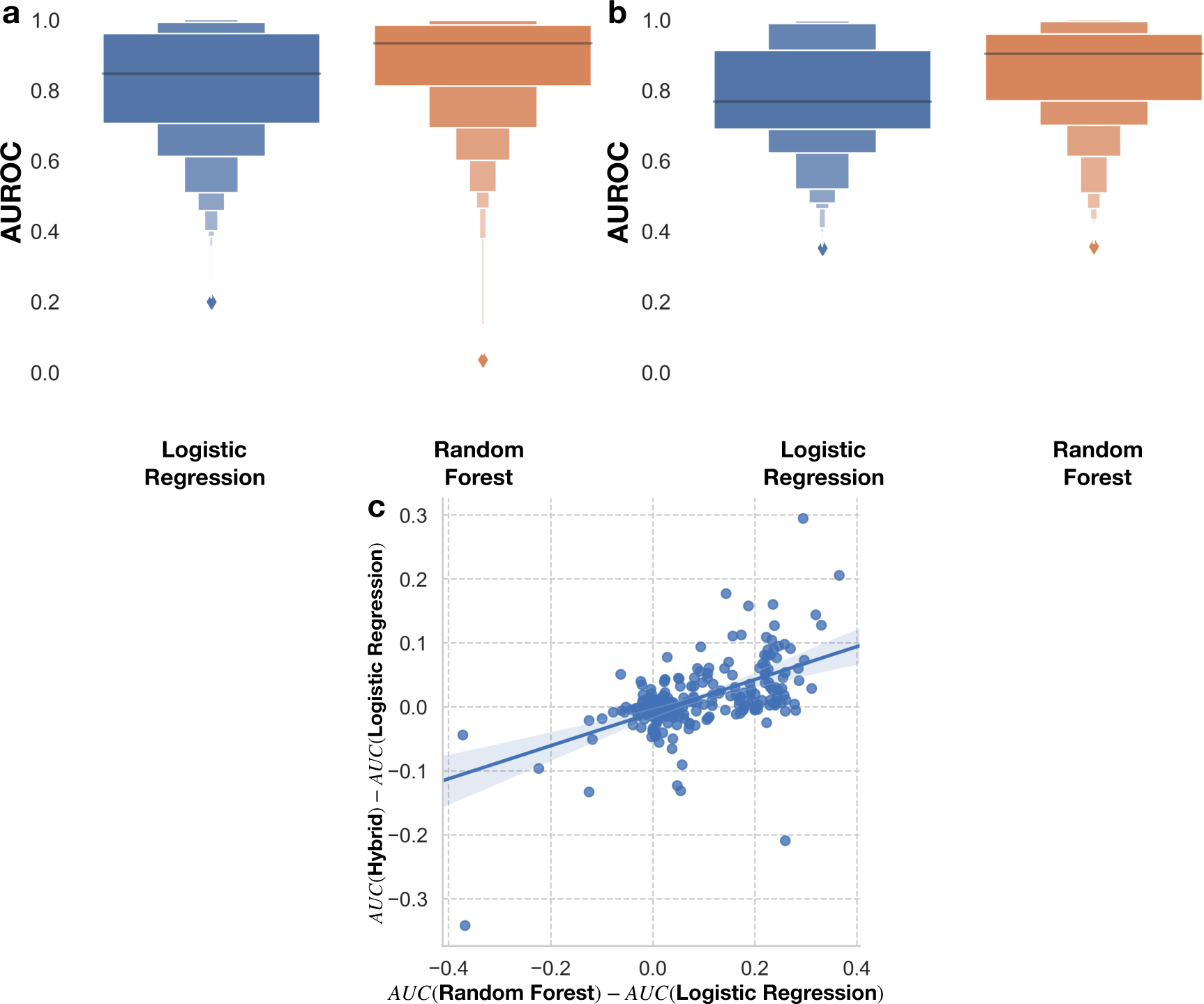
(a-b) Boxenplots of prediction performance of Logistic Regression versus Random Forest for a) original 556 datasets, b) subset 277 datasets, c) linear plot of the performance gains of hybrid approach versus the random forest approach

We then restricted range of the number of variables and samples from 5 to 110 and 50 to 2500 respectively to focus on low dimensional domains where it is more sensible to employ standard statistical modeling techniques in the biomedical space. Here, out of the remaining 277 datasets studied, the random forest model further outperformed the logistic regression models by an average 5-fold cross validated C-statistic difference of 0.077, 0.86 AUROC to 0.78 AUROC respectively (Figure 2b) (t=11.5, p=3.8e-25). At this point, we felt that it would be pertinent to study the nature of this difference through the introduction of interaction terms into the logistic regression model using our interaction extraction approach.

### Application of Interaction Terms

Out of the remaining 277 datasets, we applied the interaction terms to the model by using SHAP to add pertinent terms to the design matrix. We performed this procedure, ran a logistic regression model on the resulting design matrix, and calculated the C-statistic on a validation set for each of 5 cross validation folds, as noted in the previous section. The resulting measures of model fit were compared to the fit of the original logistic regression models (those not enhanced by the ML identified interactions) and plotted against the improvement of the random forest model over the logistic regression approach. Note that improvement of the logistic regression models may be bounded by a maximum score dictated by the maximum score attainable by any model, 100%. As seen in Figure 2c, the domain in which the random forest model is closer to the true model is when the differences in the cross-validation scores on the x-axis is greater than 0. In this case, extracting interactions proved to be beneficial when the differences in cross validation scores between the interaction and logistic regression model in the y-axis was greater than 0.

We also sought to determine how a logistic regression model augmented by interactions extracted from the machine learning model would respond when the machine learning model was closer to the true model. We observed a positive relationship between the improvement of random forest versus improvement of the interaction extraction model over the logistic regression model (m=0.26, intercept=−0.01, r=0.56, p=6.8e-24). This demonstrates the potential for utilizing the output of the machine learning model to enhance statistical models such that their predictive accuracy makes a substantial gain on that of the ML procedure, even when the random forest is (closer to) the true model. Conversely, when the random forest underperformed versus the initial logistic regression model, traditional statistical methods tended to reject the inclusion of the interactions (if the ML output even suggested them in the first place).

### Selected Biomedical Case Studies

Finally, we explore two test use cases using pertinent biomedical datasets. In the first, epistasis is studied using a model rich with interactions where the random forest model is closer to the true model and a diabetes dataset where the logistic regression model is closer to the true model to understand the nature of the extracted interactions.

### CASE STUDY 1: Interaction Extraction Helps Model Epistasis

Epistasis is defined as the interaction between multiple genes, where the complex interplay between the presence or absence of two genes in the form of alleles serves as an effect modifier to elicit the development of a complex trait or disease. Many researchers conceptualize the components of these interactions to be binary explanatory variables. The presence of these interactions can hamper the ability to uncover the true effect of the target gene in question. There are over 21,000 genes in the human genome, which can make studying all of the possible interactions between the genes computationally intractable, especially given interactions with more than two genes. Machine learning approaches such as multi-factor dimensionality reduction seek to characterize all possible pairwise interactions, but still suffer from the computational time needed to process all such interactions [28, 29].

There exist a few interacting genes that we are unable to detect, so we expect the random forest model to outperform the logistic regression model. When we train both models and test on the test set, random forest outperforms logistic regression 0.8 to 0.47, a monstrous performance gain of 0.33 (Table 1). This makes for a perfect use case from which to evaluate the ability to extract interaction terms that enhance the performance of logistic regression models.

**Table 1:** 95% confidence intervals for predictive performance (C-statistic) of Logistic Regression, Hybrid Approach, and Random Forest on held-out test set for the tasks of Epistasis and Diabetes prediction; confidence intervals obtained using 1000 sample non-parametric bootstrap

| Method | AUROC |  |
| --- | --- | --- |
|  | Epistasis | Diabetes |
| Logistic Regression | $0.473 \pm 0.0568$ | $0.831 \pm 0.059$ |
| Hybrid Approach | $0.847 \pm 0.0372$ | $0.831 \pm 0.059$ |
| Random Forest | $0.799 \pm 0.0434$ | $0.829 \pm 0.0592$ |

Inspection of SHAP summary plots reveals the salient features for each training data set for each model. It is clear that the naive logistic regression model is unable to determine which genes are interacting. The random forest plot tells another story, the SHAP plots upweight genes that appear to have interactions with other genes but does not clearly describe which genes it is interacting with. After we extract the interactions, we augmented the design matrix for the logistic regression model and estimated the corresponding logistic regression model. The results reveal that the added interactions substantially enhances the logistic regression model (AUROC 0.85) while enabling the familiar interaction-effect regression model interpretation. Inspection of the SHAP plots and exponentiated model coefficients identifies which genes were explicitly and significantly interacting with one another, and which were overall most explanatory for the complex trait being studied (Figure 3a-d). Furthermore, a study of the SHAP ranking of all of the interactions of the model yields four interactions as far stronger performers than any others (Figure 3e).

**Figure 3:**
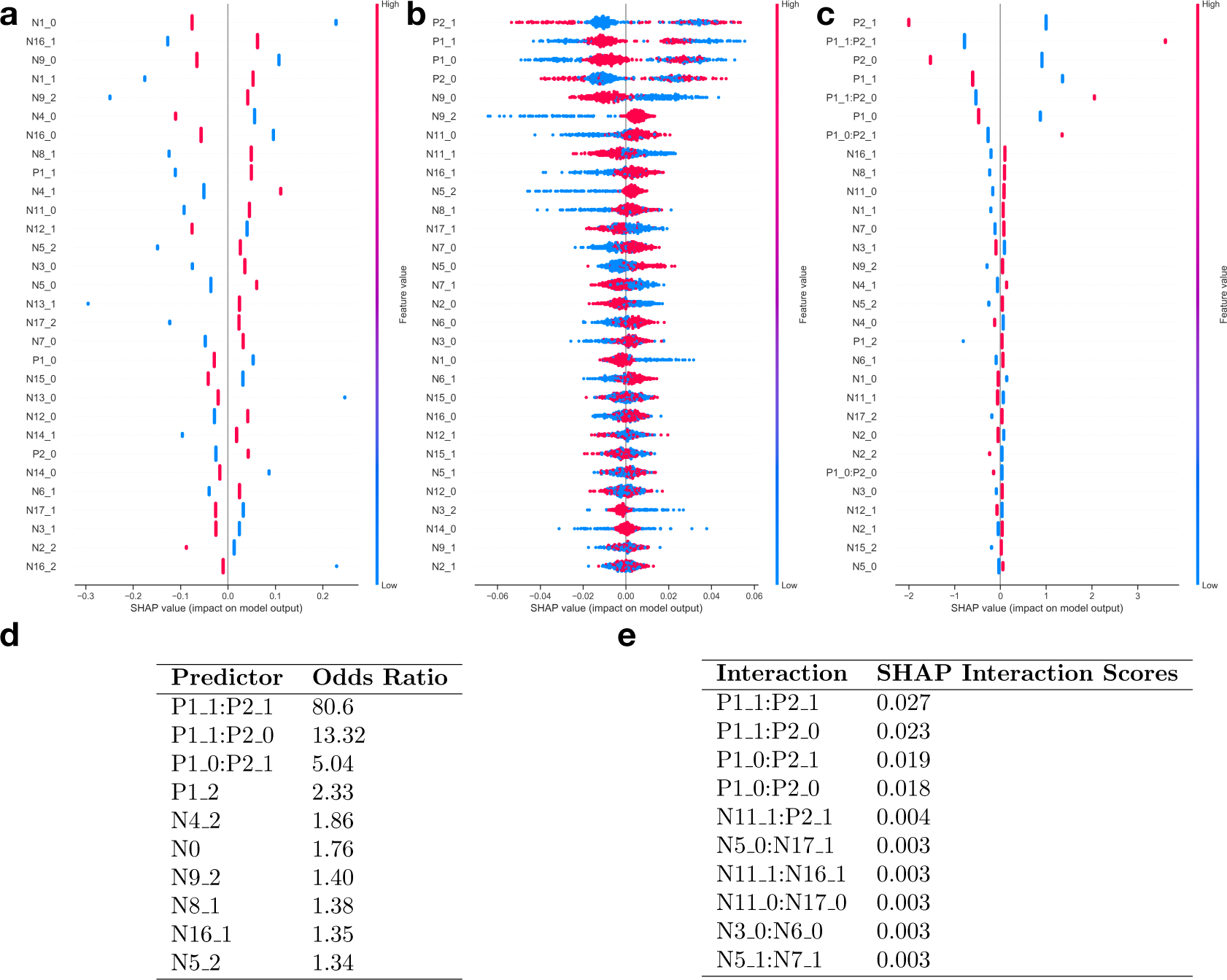
(a-c) Epistasis SHAP summary plots for: a) Logistic Regression, b) Random Forest, and c) Hybrid Approach; d) Odds-ratios for coefficients of predictors derived from hybrid logistic regression augmented by RF suggested interactions; e) SHAP scores (measures of predictor variable importance) for top 10 ranked interactions

**Figure 4:**
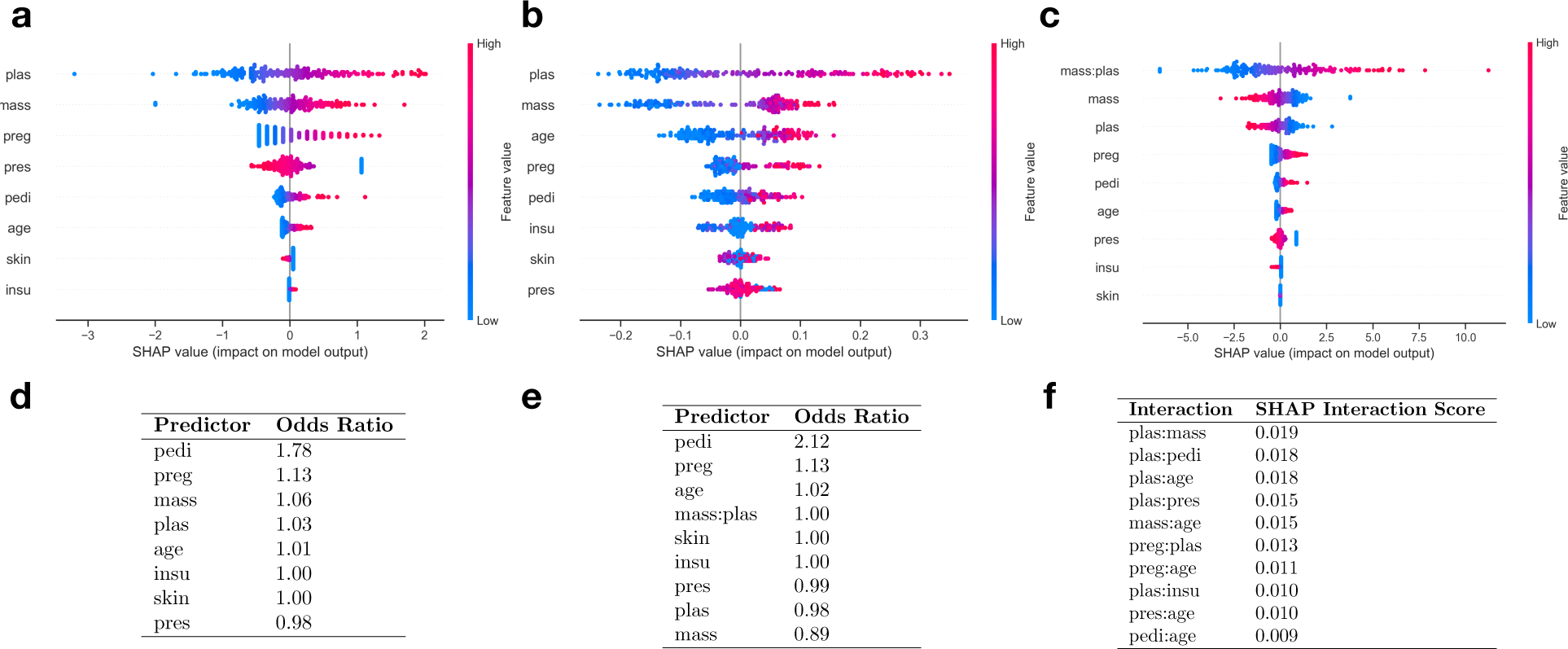
(a-c) Diabetes SHAP summary plots for: a) Logistic Regression, b) Random Forest, and c) Hybrid Approach; d) Odds-ratios of predictors derived using logistic regression prior to application of hybrid procedure; e) Odds-ratios of predictors derived from hybrid approach logistic regression coefficients; f) SHAP interaction scores for top 10 ranked interactions

Here, we have demonstrated the power to meaningfully extract interactions important to a machine learning model and add them to an underperforming logistic regression model to boost its performance to match that of the machine learning model. Thus, in this case, the hybrid approach can be viewed as outperforming either the logistic regression or Random Forest approaches alone! In our next example, we will study a case where adding interaction terms did not improve model performance.

### CASE STUDY 2: Random Forest and Logistic Regression Features Coherent for Diabetes Prediction

In the early 1900s, the diversion of irrigation systems that had once made the Pima Indians of Arizona and Mexico a prosperous agricultural community disrupted their lifestyle from one that had subsided on high carbohydrate, low fat diet to one based on a high fat diet and a more sedentary lifestyle. The Pima Indians have been willing participants in a myriad of epidemiological, clinical and genetic studies that have ultimately contributed to a better understanding of the pathogenesis and diagnosis of type II diabetes and obesity. Here, we study a dataset acquired from the National Institute of Diabetes and Digestive and Kidney Diseases and fit logistic regression and random forest models to the data to predict diabetes as a binary outcome as defined by the World Health Organization [30, 31].

The logistic regression and random forest model both exhibit similar performance (0.83 C-statistic; Table 1) while generally agreeing on which variables are the most important for the model’s decisions. After adding interactions to the logistic regression model, the performance does not change. The interaction between BMI and plasma measurements was the most significant interaction detected using SHAP (Figure 3c) and appears to be the most important variable when running it in the logistic modeling framework. This is a typical use case where extracting the interactions does not improve model performance; logistic regression is the correct model specification and it is able to strongly hint the fact to us by the non-significant effects for the interaction (guiding us to stay with the no interaction specification).

## Discussion

We have described and illustrated some of the key differences between traditional statistical modeling and machine-learning approaches, focusing the latter on a comparison to random forests. The former approach models the explanatory variables as varying linearly on the natural scale of the outcome, while the latter incorporates unguided interaction and transformation modeling. These modeling approaches are sometimes compared unfairly due to model building being inherent to ML but not to logistic regression. Yet there are many use-cases when logistic regression models outperform machine learning approaches. For future biomedical applications, the researcher should be privy to which model specification best captures the relationship between the predictor and outcome variables. In the following, we summarize the main discussion points of our work with sensible modeling recommendations:

- Machine learning approaches may be more suitable for modeling when the number of variables far outnumbers the number of features -- the high dimensionality and multi-collinearity domain. In this domain, you could use machine learning for dimensional reduction of the analysis data set [32, 33] prior to estimating logistic regression model on only the remaining terms.
- In the domain where the number of variables is similar to or less than the number of observations, testing logistic regression and random forest models on the data should give an indication of which approach is more suitable.
- If logistic regression outperforms or is on par with the random forest model, then the true model is likely to have a systematic component that is linear on the log-odds scale, and as such the results of the logistic model should be presented. There should be consistency between which features both models found to be important.
- If random forest outperforms logistic regression, the true model likely involves interactions and nonlinear variable transforms. One could apply the interaction extractions as presented in this work on the random forest model to construct an interpretable logistic regression model.
  - If the number of variables is too high, it may be computationally inefficient to extract the interaction term, nor may it improve the logistic regression model performance. In this case, the variables should be modeled using the machine learning model and explained through SHAP or alternatively the aforementioned dimensionality reduction technique should be applied.
- If interaction extraction demonstrates limited improvement, combining this approach with the variable transformer such as SAFE could potentially recover most of the performance difference.

Since variable transformations and interactions are a staple of machine learning, we envision future model building approaches to iteratively transform and form important interactions between the predictors in a model. In addition to this, we can incorporate feature grouping or feature selection processes to further eliminate issues associated with multi-collinearity and make sure that we are capturing valuable interactions. Finally, we envision that future modeling approaches may be able to hybridize machine learning methods such as Classification and Regression Trees (CART) and traditional approaches through the development of mechanisms that can automatically add smooth linear modeling specifications for the particular decision splits when a linear model can best approximate the relationship when restricted to the domain of the split. We call for derivations of more higher-level interactions and variable transformations to further enhance traditional statistical procedures. In addition, we encourage the development of methods for rigorous mathematical characterization of hybrid procedures and the conditions and specifications under which they work best.

We acknowledge that the datasets used in this study were from a machine learning benchmark repository, which means that there may be some selection bias as to the suitability of each dataset for machine learning versus traditional statistical modeling approaches. For instance, other databases such as PMLB [34] have also been developed to benchmark machine learning algorithms. Also, because the acquired datasets were based on binary prediction tasks, we did not attempt to study approaches with continuous or multinomial outcomes. Finally, our approach focused on random forest as a representative of the ML approaches. Deep learning methodologies [35] can also be used for interaction extraction, and future work could attempt to prune these neural networks until they converge on linear modeling solutions as a means to estimate complexity and number of layers of interactions needed to best model the data.

## Conclusion

We demonstrated that machine learning techniques may not always include the optimal model for a biomedical problem. In cases where they are, there may be means from which to learn important interactions from the machine learning models and apply them to enhance statistical models. Future approaches to studying biomedical problems may benefit from entwining the algorithms acquired from machine learning with the simplicity and interpretability of statistical procedures. Such work may be a step in the right direction to build trust and acceptance for machine learning into biomedical clinical settings. Our procedure and its illustration are just scratching the surface of what can be achieved by hybrid procedures. We encourage readers to consider developing or adopting more hybrid statistical – ML procedures, especially in applications of an exploratory or predictive nature for which transparent interpretation of the results is needed.

## List of Abbreviations

AUROC: Area under the receiver operating curve
CART: Classification and Regression Trees
C-statistic: Concordance statistic
CT: Computed Tomography
EHR: Electronic Health Record
MAR: Mean Absolute Error/Residual
ML: Machine Learning
PMLB: Penn Machine Learning Benchmarks
SAFE: Surrogate Assisted Feature Extraction
SAS: Statistical Analysis System
SHAP: Shapley Additive Explanations

## Acknowledgements

Not Applicable.

## Contributions

JL and JO designed the analytic plan, JL executed the analytic plan, and all authors edited and approved the final manuscript.

## Availability of data and materials

**Project Name:** InteractionTransformer

**Project home page:** https://github.com/jlevy44/InteractionTransformer

**Operating system(s):** Platform independent

**Programming language:** Python, R

**License:** MIT

The data used for evaluating the machine learning benchmark datasets are openly available for download at https://www.openml.org/. We have listed the dataset IDs from which to download these datasets in our GitHub repository, here: https://github.com/jlevy44/InteractionTransformer.

## Appendix A: Glossary of Terms

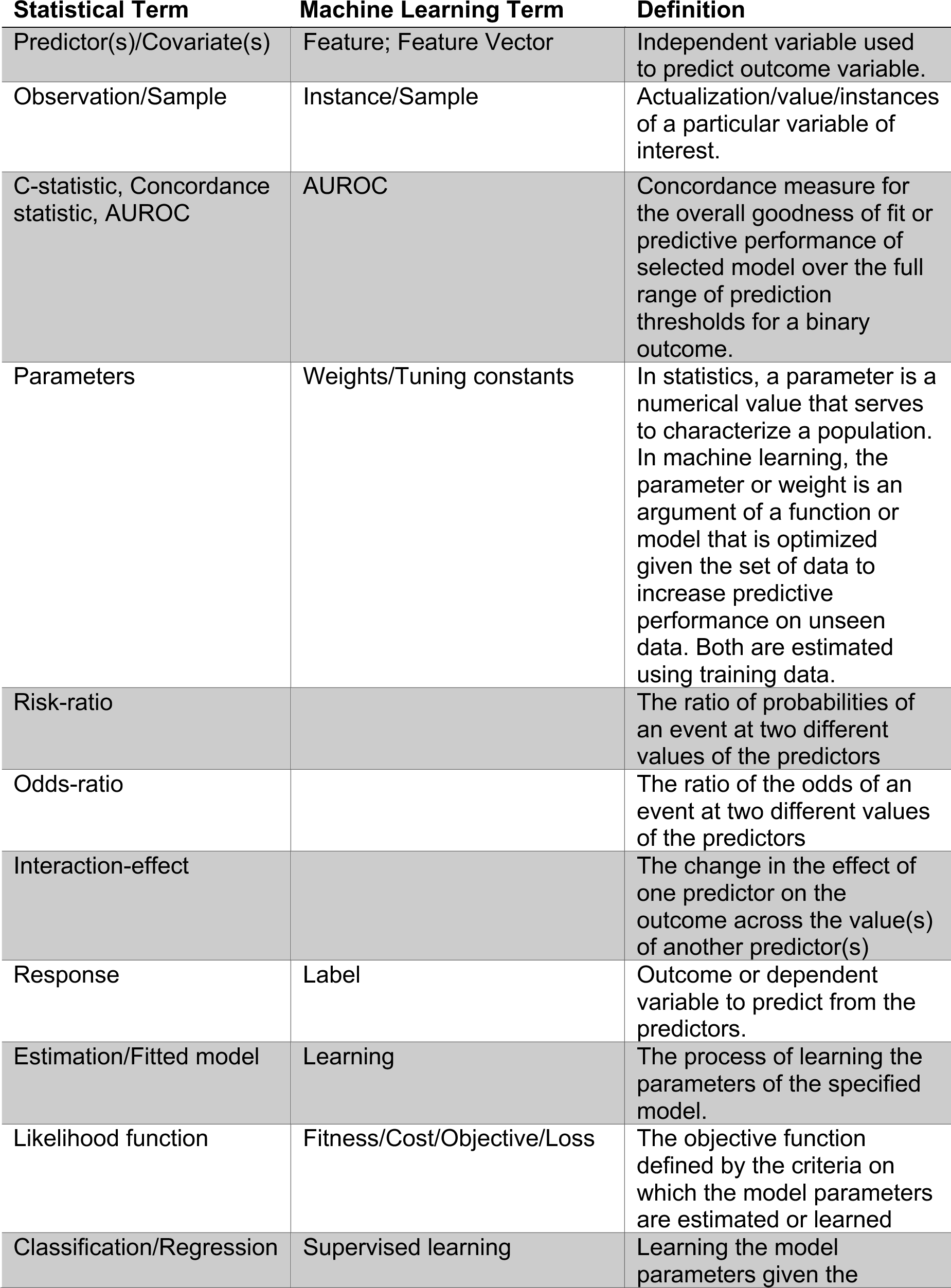
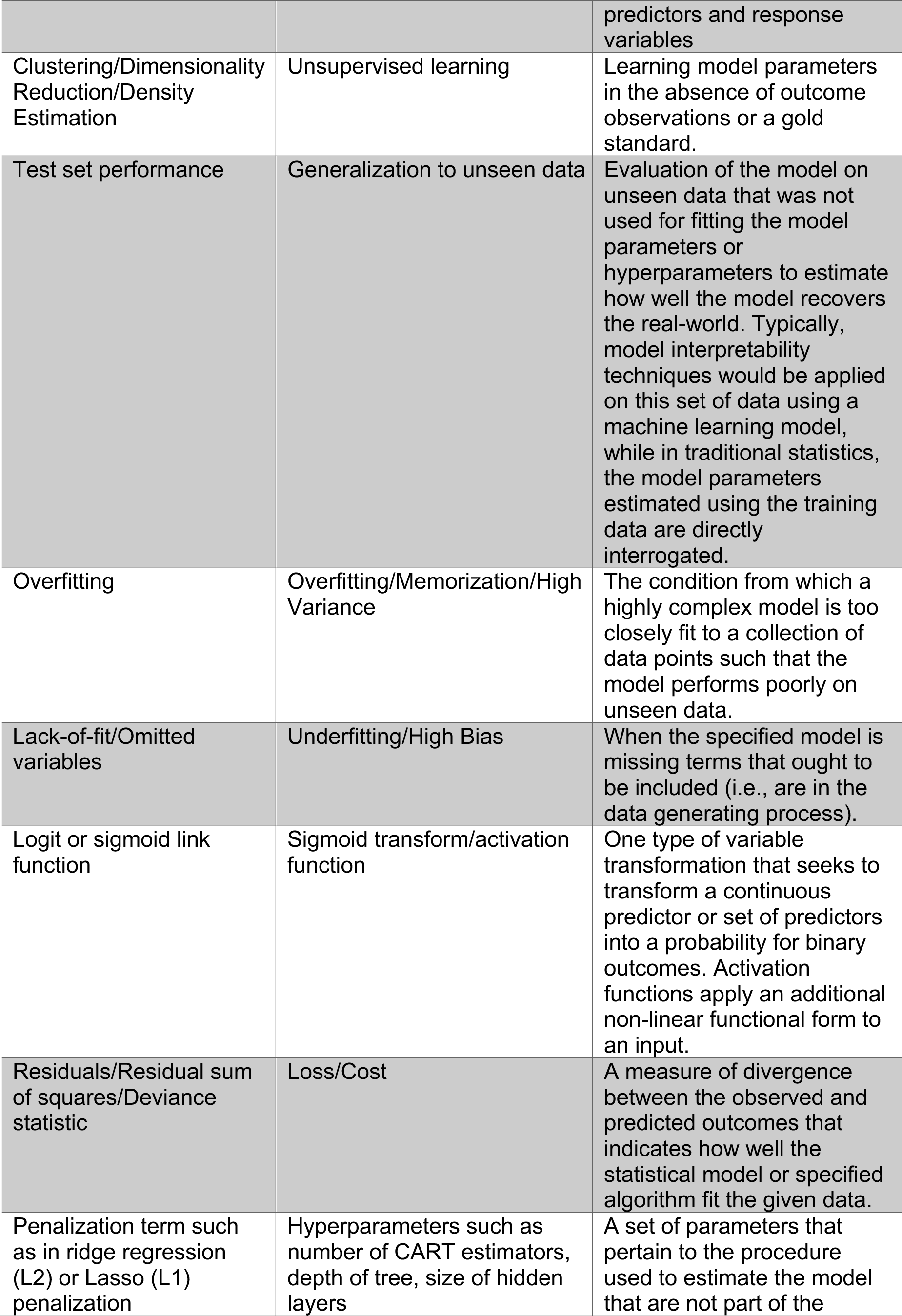
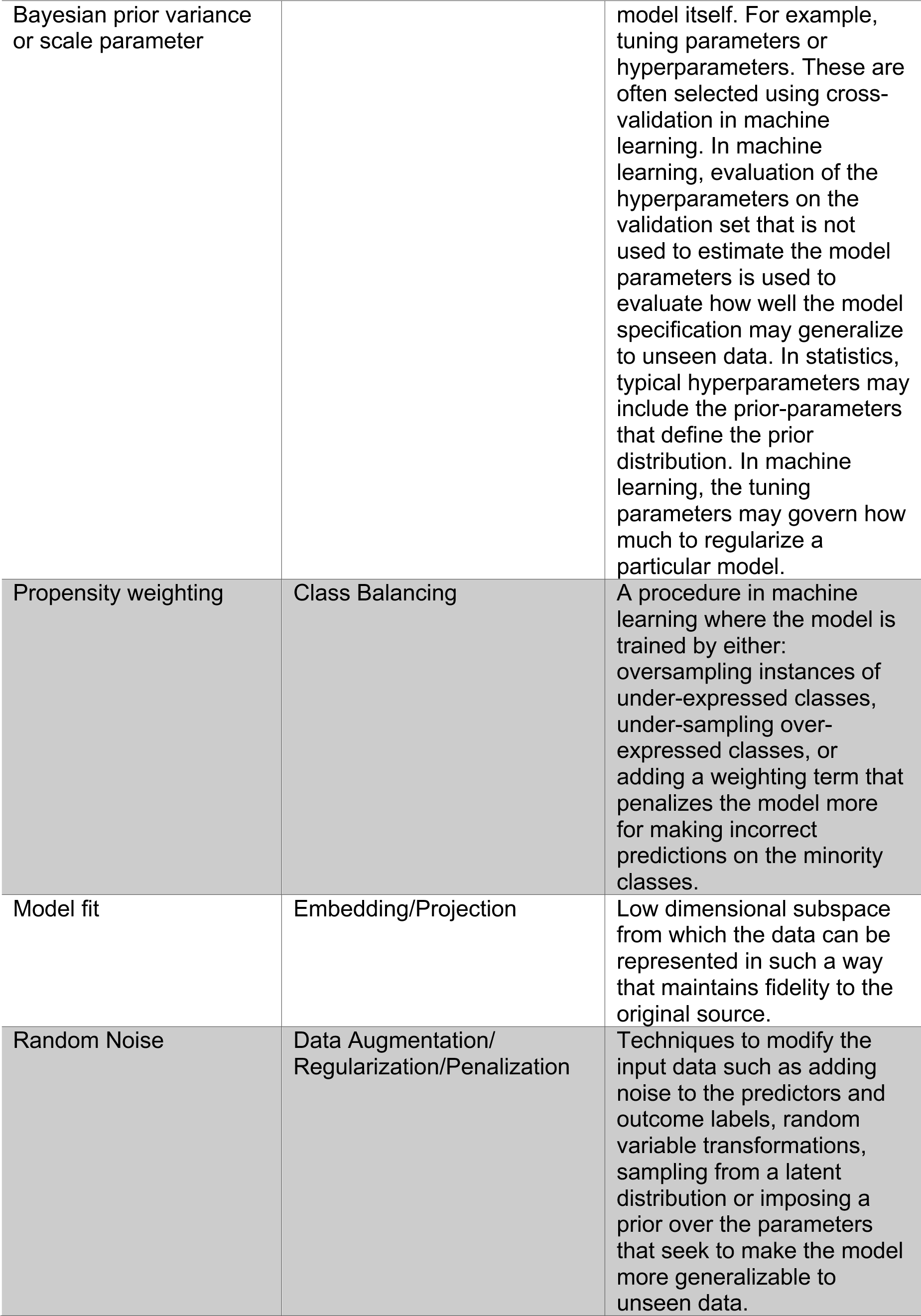
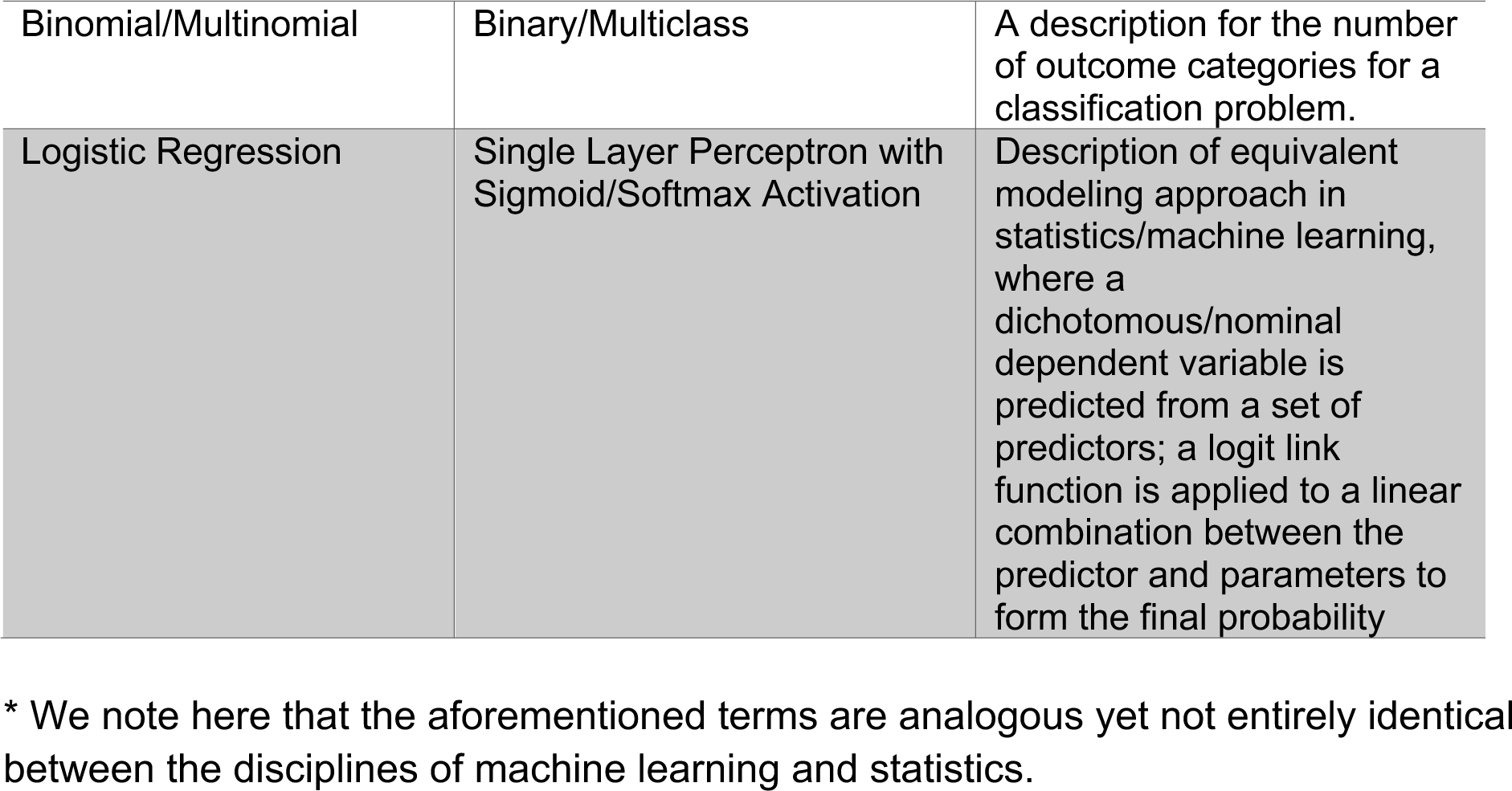

## References

1. Ching Travers, Himmelstein Daniel S., Beaulieu-Jones Brett K., Kalinin Alexandr A., Do Brian T., Way Gregory P., et al. Opportunities and obstacles for deep learning in biology and medicine. Journal of The Royal Society Interface. 2018;15:20170387.

2. Levy JJ, Titus AJ, Petersen CL, Chen Y, Salas LA, Christensen BC. MethylNet: A Modular Deep Learning Approach to Methylation Prediction. bioRxiv. 2019;:692665.

3. Chen X, Ishwaran H. Random forests for genomic data analysis. Genomics. 2012;99:323–9.

4. Jiang R, Tang W, Wu X, Fu W. A random forest approach to the detection of epistatic interactions in case-control studies. BMC Bioinformatics. 2009;10 Suppl 1:S65.

5. Cheng J-Z, Ni D, Chou Y-H, Qin J, Tiu C-M, Chang Y-C, et al. Computer-Aided Diagnosis with Deep Learning Architecture: Applications to Breast Lesions in US Images and Pulmonary Nodules in CT Scans. Sci Rep. 2016;6:1–13.

6. Coudray N, Ocampo PS, Sakellaropoulos T, Narula N, Snuderl M, Fenyö D, et al. Classification and mutation prediction from non–small cell lung cancer histopathology images using deep learning. Nat Med. 2018;24:1559–67.

7. Tang B, Li A, Li B, Wang M. CapSurv: Capsule Network for Survival Analysis With Whole Slide Pathological Images. IEEE Access. 2019;7:26022–30.

8. Ishwaran H, Kogalur UB, Blackstone EH, Lauer MS. Random survival forests. Ann Appl Stat. 2008;2:841–60.

9. Shickel B, Tighe P, Bihorac A, Rashidi P. Deep EHR: A Survey of Recent Advances in Deep Learning Techniques for Electronic Health Record (EHR) Analysis. IEEE J Biomed Health Inform. 2018;22:1589–604.

10. Couronné R, Probst P, Boulesteix A-L. Random forest versus logistic regression: a large-scale benchmark experiment. BMC Bioinformatics. 2018;19:270.

11. Christodoulou E, Ma J, Collins GS, Steyerberg EW, Verbakel JY, Van Calster B. A systematic review shows no performance benefit of machine learning over logistic regression for clinical prediction models. Journal of Clinical Epidemiology. 2019;110:12–22.

12. Khattree R, Naik DN. Applied Multivariate Statistics with SAS Software, Second Edition. 2nd edition. Cary, NC, USA: SAS Institute Inc.; 2018.

13. Johnsson T. A procedure for stepwise regression analysis. Statistical Papers. 1992;33:21–9.

14. Hocking RR. A Biometrics Invited Paper. The Analysis and Selection of Variables in Linear Regression. Biometrics. 1976;32:1–49.

15. Efroymson MA. Multiple regression analysis. Mathematical Methods for Digital Computers. 1960;:191–203.

16. Kleinbaum DG, Klein M. Introduction to Logistic Regression. In: Kleinbaum DG, Klein M, editors. Logistic Regression: A Self-Learning Text. New York, NY: Springer; 2010. p. 1–39. doi:10.1007/978-1-4419-1742-3_1.

17. Quinlan JR. Induction of decision trees. Mach Learn. 1986;1:81–106.

18. Breiman L. Random Forests. Machine Learning. 2001;45:5–32.

19. Ho TK. Random Decision Forests. In: Proceedings of the Third International Conference on Document Analysis and Recognition (Volume 1) – Volume 1. Washington, DC, USA: IEEE Computer Society; 1995. p. 278–. http://dl.acm.org/citation.cfm?id=844379.844681. Accessed 11 Apr 2019.

20. Gosiewska A, Gacek A, Lubon P, Biecek P. SAFE ML: Surrogate Assisted Feature Extraction for Model Learning. arXiv:190211035 [cs, stat]. 2019. http://arxiv.org/abs/1902.11035. Accessed 6 Nov 2019.

21. Lundberg SM, Lee S-I. A Unified Approach to Interpreting Model Predictions. In: Guyon I, Luxburg UV, Bengio S, Wallach H, Fergus R, Vishwanathan S, et al., editors. Advances in Neural Information Processing Systems 30. Curran Associates, Inc.; 2017. p. 4765–4774. http://papers.nips.cc/paper/7062-a-unified-approach-to-interpreting-model-predictions.pdf. Accessed 10 Jun 2019.

22. Lundberg SM, Erion GG, Lee S-I. Consistent Individualized Feature Attribution for Tree Ensembles. arXiv:180203888 [cs, stat]. 2019. http://arxiv.org/abs/1802.03888. Accessed 6 Nov 2019.

23. Lundberg SM, Erion G, Chen H, DeGrave A, Prutkin JM, Nair B, et al. Explainable AI for Trees: From Local Explanations to Global Understanding. arXiv:190504610 [cs, stat]. 2019. http://arxiv.org/abs/1905.04610. Accessed 6 Nov 2019.

24. Lunetta KL, Hayward LB, Segal J, Van Eerdewegh P. Screening large-scale association study data: exploiting interactions using random forests. BMC Genet. 2004;5:32.

25. Bischl B, Casalicchio G, Feurer M, Hutter F, Lang M, Mantovani RG, et al. OpenML Benchmarking Suites. arXiv:170803731 [cs, stat]. 2019. http://arxiv.org/abs/1708.03731. Accessed 6 Nov 2019.

26. Pedregosa F, Varoquaux G, Gramfort A, Michel V, Thirion B, Grisel O, et al. Scikit-learn: Machine Learning in Python. Journal of Machine Learning Research. 2011;12 Oct:2825–30.

27. Lemaître G, Nogueira F, Aridas CK. Imbalanced-learn: A Python Toolbox to Tackle the Curse of Imbalanced Datasets in Machine Learning. Journal of Machine Learning Research. 2017;18:1–5.

28. Cordell HJ. Epistasis: what it means, what it doesn’t mean, and statistical methods to detect it in humans. Hum Mol Genet. 2002;11:2463–8.

29. Sailer ZR, Harms MJ. Detecting High-Order Epistasis in Nonlinear Genotype-Phenotype Maps. Genetics. 2017;205:1079–88.

30. Schulz LO, Chaudhari LS. High-Risk Populations: The Pimas of Arizona and Mexico. Curr Obes Rep. 2015;4:92–8.

31. Acton KJ, Ríos Burrows N, Moore K, Querec L, Geiss LS, Engelgau MM. Trends in Diabetes Prevalence Among American Indian and Alaska Native Children, Adolescents, and Young Adults. Am J Public Health. 2002;92:1485–90.

32. Schölkopf B, Smola A, Müller K-R. Kernel principal component analysis. In: Gerstner W, Germond A, Hasler M, Nicoud J-D, editors. Artificial Neural Networks — ICANN’97. Berlin, Heidelberg: Springer; 1997. p. 583–8.

33. McInnes L, Healy J, Melville J. UMAP: Uniform Manifold Approximation and Projection for Dimension Reduction. arXiv:180203426 [cs, stat]. 2018. http://arxiv.org/abs/1802.03426. Accessed 5 Mar 2019.

34. Olson RS, La Cava W, Orzechowski P, Urbanowicz RJ, Moore JH. PMLB: a large benchmark suite for machine learning evaluation and comparison. BioData Mining. 2017;10:36.

35. LeCun Y, Bengio Y, Hinton G. Deep learning. Nature. 2015;521:436–44.

